# Towards a more robust non-invasive assessment of functional connectivity

**DOI:** 10.1101/2023.02.06.527279

**Authors:** Britta U. Westner, Jan Kujala, Joachim Gross, Jan-Mathijs Schoffelen

## Abstract

Non-invasive evaluation of functional connectivity, based on source-reconstructed estimates of phase-difference-based metrics, is notoriously non-robust. This is due to a combination of factors, ranging from a misspecification of seed regions to suboptimal baseline assumptions, and residual signal leakage. In this work, we propose a new analysis scheme of source level phase-difference-based connectivity, which is aimed at optimizing the detection of interacting brain regions. Our approach is based on the combined use of sensor subsampling and dual-source beamformer estimation of all-to-all connectivity on a prespecified dipolar grid. First, a pairwise two-dipole model, to account for reciprocal leakage in the estimation of the localized signals, allows for a usable approximation of the pairwise bias in connectivity due to residual leakage of ‘third party’ noise. Secondly, using sensor array subsampling, the recreation of multiple connectivity maps using different subsets of sensors allows for the identification of consistent spatially localized peaks in the 6-dimensional connectivity maps, indicative of true brain region interactions. These steps are combined with the subtraction of null coherence estimates to obtain the final coherence maps. With extensive simulations, we compared different analysis schemes for their detection rate of connected dipoles, as a function of signal-to-noise ratio, phase difference and connection strength. We demonstrate superiority of the proposed analysis scheme in comparison to single-dipole models, or an approach that discards the zero phase difference component of the connectivity. We conclude that the proposed pipeline allows for a more robust identification of functional connectivity in experimental data, opening up new possibilities to study brain networks with mechanistically inspired connectivity measures in cognition and in the clinic.

## 1 Introduction

The brain is considered to operate as a network of interacting, functionally specialized regions. The development and application of analysis tools to probe those interactions in the healthy human brain from non-invasive electrophysiological measurements has been an active area of research in the past few decades. Part of that work is grounded in the notion that interregional interactions may be reflected by statistical dependencies between band-limited signal components that can be picked up from locally activated brain areas. One way to quantify this so-called functional connectivity is to estimate some measure of relative phase consistency or phase synchrony (Varela et al., 2001), for instance using the coherence coefficient, or a derived metric (Bastos and Schoffelen, 2016). From a mechanistic point of view it has been hypothesized that consistent phase differences of oscillatory processes facilitate neuronal interactions by virtue of a mutual temporal alignment of cycles of increased neuronal excitability (Fries, 2005, 2015; Bonnefond et al., 2017). In sum, connectivity estimates based on phase synchrony are a valuable metric in cognitive neuroscience.

It is commonly agreed that, for interpretability, connectivity estimates should be assessed at the source level. This is because connectivity estimates are invariably confounded by spatial leakage (Schoffelen and Gross, 2009). Promising work from the early 2000s developed (Gross et al., 2001) and successfully applied (e.g., Pollok et al., 2005; Schoffelen et al., 2005) the Dynamic Imaging of Coherent Sources (DICS) technique, a frequency domain version of a beamformer for source reconstruction, to identify networks of phase synchronized brain regions based on the strong physiological periodicities during smooth finger movements in healthy participants. Further studies focused on synchrony at the frequency of Parkinsonian or essential tremor in clinical populations (Timmermann et al., 2003; Pollok et al., 2004). In the decades following this early work, the research community has also started studying envelope correlations of band-limited signals instead of phase synchrony. This latter metric has been successfully used to identify properties of networks predominantly during the brain’s resting state, yielding a body of literature with well interpretable and consistent findings (Brookes et al., 2011; Hipp et al., 2012; Baker et al., 2014; Colclough et al., 2016; de Pasquale et al., 2016). Despite ongoing methodological work to improve source reconstruction (Dalal et al., 2006; Woolrich et al., 2011; Hillebrand et al., 2012; Nunes et al., 2020; Kuznetsova et al., 2021) and novel phase synchrony based connectivity metrics (Aviyente et al., 2011; Vinck et al., 2011; Ghanbari and Moradi, 2020), neuroscientific findings employing phase synchrony seem to be more scarce and less consistent (Colclough et al., 2016; O’Neill et al., 2018).

Assuming that metrics based on phase differences tap into fundamental mechanisms of brain organization and communication (Fries, 2005, 2015; Bonnefond et al., 2017), then why is it seemingly so difficult to find converging evidence across studies? One reason for this might be that the methodological adversities are larger than commonly assumed (Bastos and Schoffelen, 2016; Palva et al., 2018; He et al., 2019). One of those difficulties is spatial leakage, both from second party and third party sources, which renders the lower bound of the true connectivity unknown. Proposed techniques for leakage correction, on the other hand, might be too aggressive and also compromise or even remove the signal of interest. Furthermore, data quality might further impede the reliable estimation of phase difference: low signal-to-noise ratio (SNR) might hinder the reliable identification of seed regions of interest, while SNR differences across conditions occlude the interpretation of connectivity, since the estimation of phase-based connectivity measures is sensitive to SNR changes.

In this paper, we propose a new method that tackles these problems. We propose to address the issue of suboptimal region of interest (ROI) or seed selection through consideration of the full 6-dimensional all-to-all connectivity source space, using a two-dipole constraint beamformer. We further propose an estimation of the null coherence which approximates the bias in the coherence estimate and can be used to correct the output. Finally, we reduce estimation bias by aggregating over the results of many source reconstructions using sensor array subsampling, thereby creating a more stable and robust estimate.

In the following, we will first introduce beamforming for source reconstruction and explain the problem of spatial leakage in more detail. Then we will line out the components of our proposed beamforming approach.

### 1.1 Beamformers for source reconstruction

Non-invasive electrophysiological measurements (electric potential differences for electroencephalography (EEG) or magnetic fields (gradients) for magnetoencephalography (MEG)) reflect a mixture of the temporal activation profiles from neural and non-neural sources. To disentangle the individual contributions of each of those sources to the spatiotemporal mixture in the observed signals, source reconstruction techniques can be applied. These techniques have developed into a valuable tool for the analysis of non-invasive electrophysiological signals obtained during cognitive experiments. Solving the so-called inverse problem by combining a forward model with additional assumptions, source reconstruction techniques aim to build models of the spatiotemporal characteristics of the neural generators that underlie the measured signals, unmixing the observed channel-level data. The biologically plausible forward model (or gain matrix) describes the spatial distribution of the observed signals, typically for a set of equivalent current dipole sources. The additional model assumptions are necessary to constrain the number of solutions to the inverse problem, which in principle are unlimited. Adaptive beamformers are a class of source reconstruction techniques that do not a priori make explicit assumptions with respect to the number or location of active sources, but rather assume the underlying sources to be temporally uncorrelated. Usually, for each of a set of predefined source locations, a spatial filter is constructed under two constraints: 1) a unit gain constraint, which means that it should pass on all of the activity that originates from that specific location, and 2) a minimum variance constraint, which minimizes the variance of the reconstructed actibty at each location. Mathematically, this linearly constrained minimum variance (LCMV; Van Veen et al., 1997) spatial filter is computed as follows:

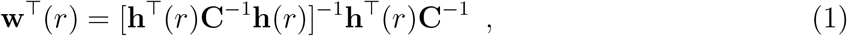

where **w**(*r*) is the spatial filter at source location *r* and ⊤ refers to the transpose operation. **h**(*r*) is the source location-specific gain vector (which can be thought of as a spatial fingerprint), and **C**^*−*1^ is the mathematical inverse of the channel covariance matrix. As an alternative to the channel level covariance matrix, one can use a complex-valued cross-spectral density (CSD) matrix, based on the channel Fourier coefficients for a given frequency bin, resulting in the DICS algorithm (Gross et al., 2001).

Beamformers have gained prominence as one of the most popular source reconstruction techniques because they typically provide relatively robust reconstructions of neural activity without the need of sophisticated parameter tweaking (Westner et al., 2022). However, some limiting factors exist with respect to functional connectivity. In the following, we will present the typical distortions when source reconstructing functional connectivity, as well as our approach to mitigate these.

### 1.2 The effect of signal leakage on source connectivity estimates

In the context of connectivity estimation, an important concept is that of signal leakage. This refers to the fact that each location’s estimated activity reflects an unknown mixture of the true activity at this location and signal contributions from distant noise sources of both neural and non-neural origin. Mathematically, this can be shown as detailed below.

Considering the generative model of the sensor-level data, the sensor signals reflect a summation of the underlying source signals, each multiplied by their spatial fingerprint:

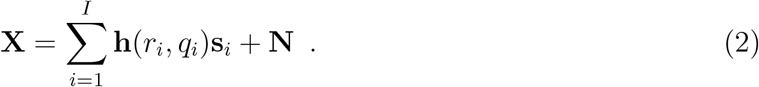

Here, **X** is a number-of-channels by number-of-observations matrix with complex-valued Fourier coefficients, **h** is the gain vector for a dipolar source at location *r*_*i*_ and with orientation *q*_*i*_, and **s**_*i*_ is a 1 by number-of-observations source activity vector, here assumed to be complex-valued, *i*.*e*., to reflect both amplitude and phase for the observations. **N** is a number-of-channels by number-of-observations matrix, reflecting the non-brain noise in the measured data.

Assume that we have computed a pair of spatial filters, **w**_1_ and **w**_2_, and we use these spatial filters to compute an estimate of the source level Fourier coefficients: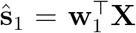, and 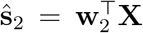. From these estimates one can compute a connectivity metric, for instance the coherence coefficient, for this dipole pair:

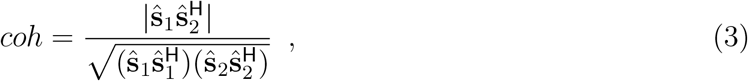

where **H** denotes the conjugate transposition. Note that for simplicity of notation, we omit the scaling with the number of observations, which drops out of the equation anyhow. We also note that a non-zero numerator in the equation above suggests linear dependence between the estimated sources 1 and 2. Below, we inspect this quantity, *i*.*e*., the cross-spectral density estimate between the two sources, in more detail.

For the given pair of dipoles, and considering the data model **X** = **h**_1_**s**_1_ + **h**_2_**s**_2_ + **N** with **N** now reflecting all signal contributions to the observed data that are not originating from the two dipole pairs-of-interest, we can express the cross-spectral density estimate between the two dipoles as:

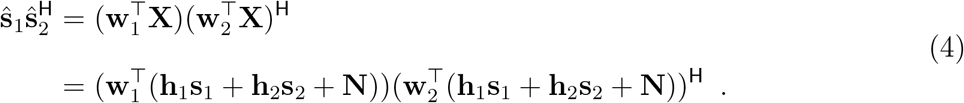

Introducing *g*_*ij*_ as a scalar value that results from computing the inner product between spatial filter **w**_*i*_ and gain vector **h**_*j*_ and which reflects the filter’s gain at location *i* for a source originating from location *j*, we obtain:

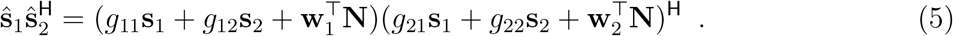

When using an inverse algorithm with a typical unit-gain constraint,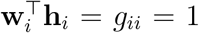, the above further reduces to:

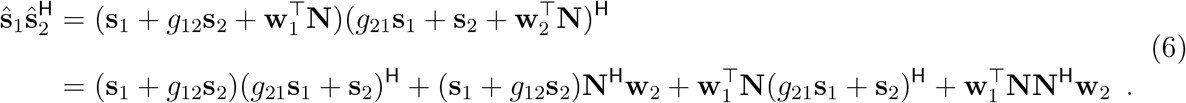

In other words, for a given dipole pair, the estimated cross-spectral density between two sources does not only depend on the sources’ true cross-spectral density, but is also affected by:

1. signal leakage from the other dipole-of-interest, specifically when *g*_12_ and *g*_21_ are non-negligible, *cf*. the leftmost term in the above equation
2. the interaction between the noise, projected through the spatial filter, and the sources’ activity, *cf*. the middle two terms in the above equation
3. the interaction between the projected noise at the location of the dipoles, *cf*. the rightmost term in the above equation

Note, that the above reasoning is independent of the exact inverse algorithm used. The different types of leakage will also affect the estimates of the individual dipoles’ power. Leakage will always cause misestimation of metrics that are derived from the estimated source level quantities. This also applies to spatial maps of connectivity, which are typically constructed using a limited number of predefined seed dipole locations. Local maxima in these spatial maps (which are either expressed as a difference between two experimental conditions or in relation to a baseline) are then interpreted as regions that are functionally connected to the seed dipole. Irrespective of the specific connectivity metric used, spatial structure in these maps due to leakage may lead to inference of false positive connections. Furthermore, true connections may be missed altogether, if the seed dipoles have been misspecified by the researcher.

### 1.3 Alleviating the effect of leakage

In order to address some of the problems associated with leakage, it has been proposed to use connectivity metrics that disregard the interaction along the real-valued axis (*e*.*g*., the imaginary part of coherency (Nolte et al., 2004) or the multivariate interaction measure (MIM, Ewald et al., 2012)), or to remove the instantaneous leakage originating from one or more dipoles prior to estimating the connectivity on the residuals (Brookes et al., 2012; Hipp et al., 2012; Colclough et al., 2015; Wens et al., 2015). Although these adjustments avoid an overinterpretation of leakage-affected findings, the sensitivity to true signal interactions at small phase differences is diminished. In addition, these leakage correction schemes do not eliminate the necessity to statistically evaluate the estimated connectivity against a well-defined null hypothesis. This step is usually not straightforward since an appropriate baseline is not available: either because of differences in the signal specific to condition or subject group (see *e.g.*, Bastos and Schoffelen, 2016), or because of the absence of a baseline condition altogether (e.g., in single group resting-state studies). Finally, in a context where seed-based connectivity maps are evaluated, there is no guarantee that the seed regions have been appropriately specified.

In this work, we propose an analysis scheme of source level connectivity (here expressed as the coherence coefficient), accounting as much as possible for the effects of leakage but without a reduction in sensitivity for true interactions at small phase differences. Moreover, we will derive estimates of a usable lower bound of the estimated coherence between dipole pairs, which can be used as a correction to more accurately evaluate spatial maps of connectivity, thus avoiding the issues associated with inappropriate or absent baseline conditions. Using extensive simulations, we show superiority of our analysis scheme in comparison to other approaches.

### 1.4 Proposed analysis approach

The analysis approach we outline in this paper consists of several elements: We make use of a two-dipole constraint beamformer (Dalal et al., 2006; Brookes et al., 2007; Schoffelen et al., 2008; Moiseev et al., 2011), approximate and correct for the estimated bias due to noise leakage, and embed the approach in a sensor array subsampling scheme. Below, we will discuss all those elements in more detail.

#### Two-dipole constraint beamformer and bias estimate

The approach is based on an all-to-all approach, where coherence is estimated between all pairs of beamformer reconstructed dipoles defined on an evenly spaced 3-dimensional grid, covering the entire brain. Using a two-dipole constraint in a beamformer formulation, we compute pairwise spatial filters that are not corrupted by zero lag correlations for the dipole pair under consideration. A beamformer with two dipoles in its spatial passband has an identity gain constraint:

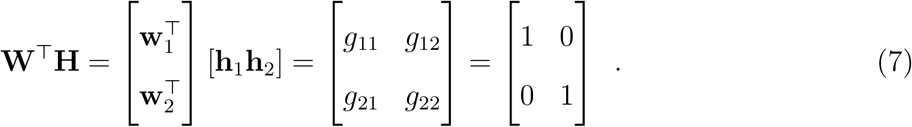

As a consequence, the equation that expresses the estimated pairwise dipolar cross-spectral density reduces to the below equation, in analogy of the model formulation as used in the previous section:

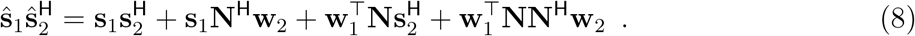

Under the assumptions that the cross terms between the noise (*i*.*e*., the part of the measured signal that does not originate from the locations of interest) and the considered sources is negligible, and if the noise covariance is assumed spatially white (a scaled identity matrix), then the above equation reduces to:

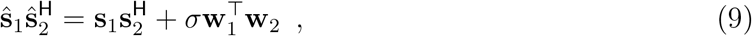

where *σ* is a scalar parameter. Under the described (yet most likely often violated) assumptions, if the true interaction strength between the two dipoles is zero, 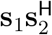 in eq. 9 will be zero. Thus, the estimated cross-spectral density between the two sources may be approximated with a scaled version of the spatial filters’ inner product, 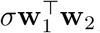, where the scaling parameter is a function of the pair of source locations. From this follows, that the scaled spatial filter inner product can be used as an approximation of the bias in estimated connectivity under the assumption of no interaction between the considered sources. Pragmatically, we propose to assume the scaling parameter to be fixed for a given seed dipole (*i*.*e*., keeping one of the dipoles in the pair fixed, and scanning through the dipole grid for the other dipole of the pair), and thus allow for its estimation by fitting a regression line through a two-dimensional point cloud, which reflects on one dimension the estimated cross-spectral density between the seed dipole and all other dipoles, and on the other dimension the inner product between the seed dipole’s spatial filter and the other dipoles’ spatial filters. Repeating this fitting procedure for all dipoles and normalizing by the product of the estimated power yields a 6-dimensional volume of null-coherence estimates, which can be used to subtract from the estimated coherence (*cf*. Fig. 1B). The resulting 6-dimensional differential map can subsequently be post-processed (*e*.*g*., thresholded) and inspected for local maxima, which might be indicative of truly interacting dipoles.

**Figure 1:**
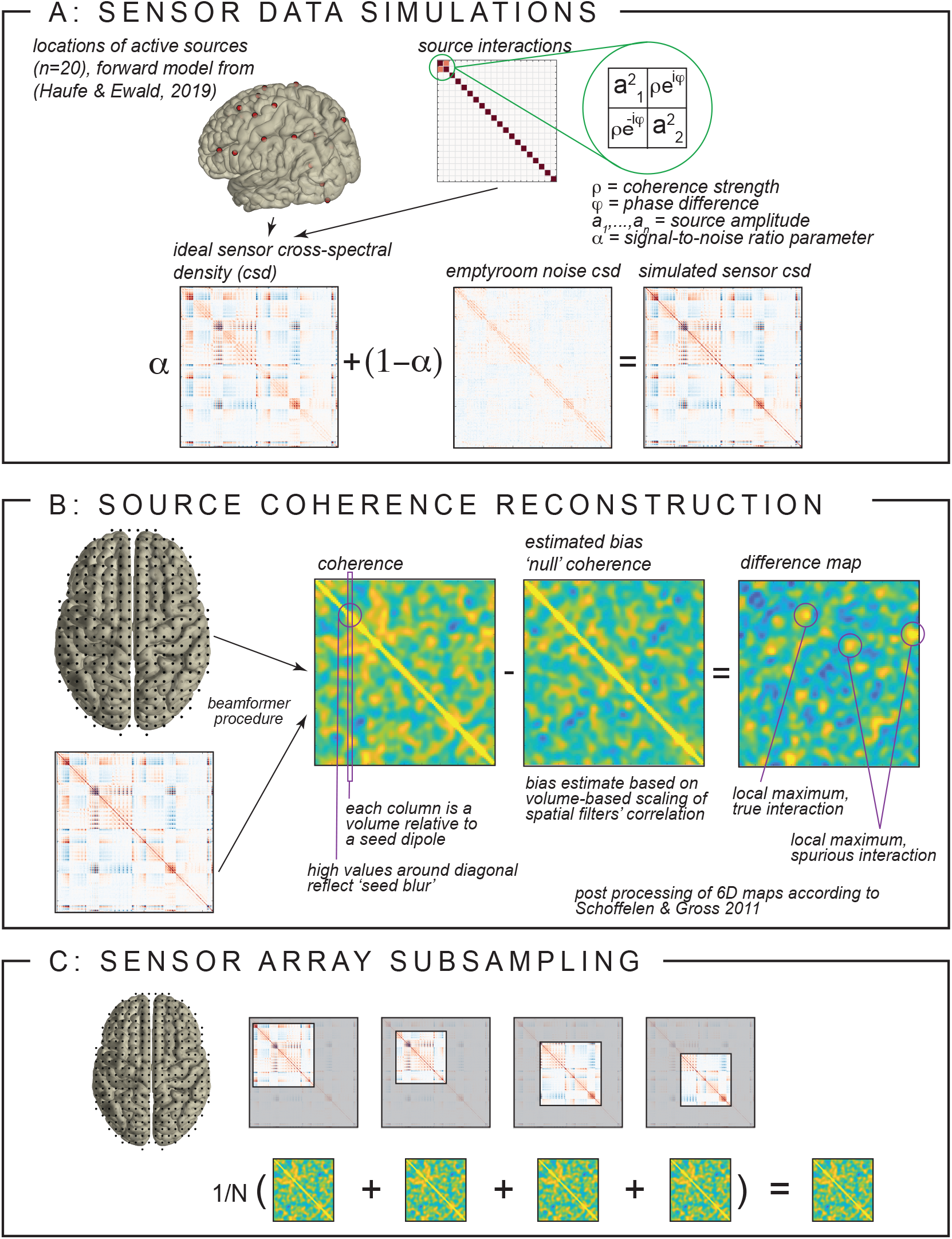
Simulation and algorithm details. **A** Setup of sensor data simulation, illustrating the interacting and non-interacting sources and signal-to-noise ratio. **B** Estimation of the null coherence across space and the computation of the difference maps. **C** Illustration of the sensor array subsampling procedure with a varying number of sensors among realizations.

#### Array subsampling

As will become clear below, these difference maps may still be spatially noisy, resulting in false positive connections (*i*.*e*., local maxima that do not reflect interacting dipoles), and true connections being missed (*i*.*e*., reconstructed connectivity between locations close to interacting dipoles not presenting as local maxima). To further reduce the spatial noise in the images, we propose a sensor array subsampling approach (Schoffelen et al., 2012; Westner et al., 2015; Westner, 2017). We estimate the 6-dimensional differential connectivity map multiple times, each time using a different random subset between 50 and 150 sensors for the reconstruction (*cf*. Fig. 1C). Although the reconstructions with fewer sensors may have a compromised spatial resolution, the spatial noise will be variable across reconstructions and thus average out, while the true interactions will show up more consistently. This scheme is akin to the idea of Ensemble Methods in machine learning, where the aggregation of many weak learners leads to a strong model with reduced variance (Breiman, 1996).

## 2 Methods

All simulations and reconstructions were performed in MATLAB (version 2021b) on a Linux operated High Performance Compute cluster, using FieldTrip (Oostenveld et al., 2011) and custom written code.

### 2.1 MEG sensor data simulations

MEG sensor space complex-valued data matrices were simulated from source space activity, based on a 275-channel axial gradiometer CTF system, as a combination of an ‘ideal’ sensor-level signal data matrix **X**_*s*_ and a noise data matrix **X**_*n*_. These *N*_sensor_ *× N*_observation_ matrices reflect the Fourier coefficients (*i*.*e*., amplitude and phase information) computed for a given frequency. For the noise matrix we used a multitaper spectral estimate of a frequency band centered around 10 Hz from a 50 second empty room measurement, recorded at the Donders Centre for Cognitive Neuroimaging. The empty room data were segmented into 1 second epochs and spectrally transformed, using a multitaper smoothing parameter of *±* 4 Hz (7 tapers per segment), which resulted in a 268 *×* 350 noise matrix. The number 268 reflects the number of active SQUIDs at the time of the empty room measurement, 350 the number of observations (*N*_epochs_ *× N*_tapers_). The signal data matrix was constructed using the generative model **X**_*s*_ = **HS**, using a precomputed forward model **H** (see below), and an *N*_source_ *×N*_observation_ matrix **S**. The source signals were simulated using MATLAB’s mvnrnd function, generating multivariate Gaussian data, with a mean of 0, and a parametrized covariance (cross-spectral density) matrix, defined as:

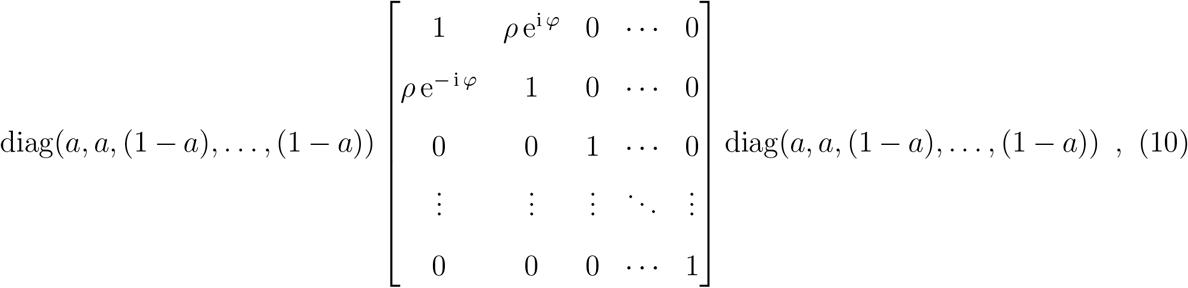

where *ρ* reflects the intended coherence coefficient between the first 2 sources and *φ* reflects the phase difference. *a* reflects a relative amplitude parameter, determining the relative amplitude of the connected dipoles in relation to the other active sources such that the relative strength between connected dipoles and active sources can be computed as *a/*(1 *− a*), *i*.*e*., a relative amplitude of 0.8 yields the connected dipoles being four times stronger than the other active sources. The procedure for simulating the sensor space data is illustrated in Figure 1A. For gain matrix **H**, we used a precomputed forward model, as described in Haufe and Ewald (2019) and the Biomag conference 2016 data analysis challenge (see https://bbci.de/supplementary/EEGconnectivity/BBCB.html). Briefly, source locations were sampled from a cortical segmentation based triangulated mesh, originally consisting of 2004 positions. A three-shell boundary element method (BEM) had been used to compute the forward solution for the 2004 dipoles with an orientation perpendicular to the cortical sheet, using Brainstorm (Tadel et al., 2011). For the simulations presented here, sets of 20 positions were randomly selected from a subset of 820 positions. This subset was created based on the norm of the gain vectors for the orientation-constrained dipoles placed at those positions: We excluded candidate locations for which the sensor array was relatively insensitive, *e*.*g*., deep dipoles in the midline, or dipoles with an unfavorable orientation. The matrices **X**_*n*_ and **X**_*s*_ were scaled with the Frobenius norm of their respective cross-spectral densities (**XX**^H^) and linearly combined using:

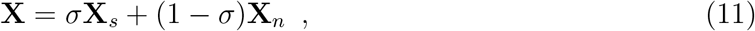

where *σ* is a parameter that determines the signal-to-noise ratio. Table 1 summarizes the relevant parameters for the simulations and the values used to explore the different reconstruction approaches.

**Table 1:** Simulation parameters. Values marked with asterisks denote values for which the outcomes are reported in the supplementary material.

| parameter | values |
| --- | --- |
| # of active sources | 20 in 100 different configurations |
| $\sigma$, signal-to-sensor-noise | 0.5*, 0.6 |
| $a$, amplitude relation | 0.5*, 0.7, 0.8* |
| $\rho$, coherence coefficient | 0, 0.2, 0.3, 0.4, 0.5, 0.6, 0.7, 0.8 |
| $\pi$, phase difference | 0, $(2/17)\pi$, $(4/17)\pi$, $(8/17)\pi$, $(16/17)\pi$ |

### 2.2 Beamformer source reconstruction and coherence estimation

For source reconstruction we used a forward model defined on a regularly spaced 3-dimensional dipole grid (with a spacing of 8 mm). The brain compartment of this grid consisted of 4416 dipoles and was defined by the same anatomical MRI as the one used for the simulations’ forward model. For the reconstructions’ forward model, we used a single shell model as implemented in FieldTrip (Nolte, 2003). Our detailed analysis required the computation of 4416^2^ pairs of spatial filters for many iterations of sensor array subsamples (we used 100 subsamples per simulation) over 8000 parameter combinations. Thus, we had to estimate over 15 trillion spatial filters in total. We wrote custom code for the efficient computation of the spatial filters and the derived coherence. All beamformers were computed with FieldTrip’s fixedori constraint, which computes a fixed orientation forward model for each dipole, based on the maximization of the beamformer’s output power (Sekihara and Nagarajan, 2008).

## 3 Results

### 3.1 Illustrative example and null coherence estimation

This section illustrates our proposed approach. Figure 2A shows the spatial configuration of one instantiation of the simulation, where 20 dipole locations were randomly selected to reflect the active sources. Two of these sources (the bigger, orange dots in the figure, here denoted as a medial superior frontal (MSF) and left occipital (LO) source) reflect the interacting dipoles. To illustrate the potential issues related to spatial leakage, we start by investigating different seedbased maps. In this example, we simulated the interaction to be at a phase difference of (8*/*17)*π* and the coherence strength to be 0.5. For illustration purposes we computed these seed-based results on a 4 mm grid, but for the reconstruction of all pairwise interactions we used an 8 mm grid. For this example, we also simulated data using identical source parameters as for the above simulations, apart from the coherence strength, which was set to zero. This simulation was intended to reflect a perfect baseline, where everything except the interaction strength was kept constant. We start the illustration using a traditional single dipole beamformer. Figure 2B shows the seed-based estimate of coherence for truly interacting sources, using as a seed the grid position that was closest to the MSF source (indicated with a white square). Figure 2C shows an estimate of the coherence for the scenario in which the dipoles were not connected. Both estimates are dominated by the well-known seed blur, but also show a small local maximum in the vicinity of the LO source (indicated with a red square) for the case of the true connectivity (Fig. 2B). The difference image (Fig. 2E) shows an effective suppression of the leakage close to the seed location. Yet, there is considerable spatial structure in the residual image, and although there is a local maximum in the vicinity of the LO source, there are also other maxima in this image that may be mistaken for interacting sources. In many practical situations, an appropriate baseline condition is not available. This motivated us to estimate the ‘null’ coherence based on the scaled inner product of the spatial filters (as described above), assuming this scaling to be fixed for a given seed dipole, and the noise to be spatially white and uncorrelated with the sources. Figure 2D shows the computed null coherence for our illustrative example. The null coherence map shows considerably more structure than the baseline condition in Figure 2C, and thus, the difference map between the coherence and null coherence (Fig. 2F) also exhibits more substantial structure than the difference map with the baseline condition (Fig. 2E). Specifically, the seed blur does not seem to be very well accounted for given the difference map’s local maximum in the vicinity of the MSF seed region. However, the local maximum in the vicinity of the LO source is more prominent in this figure than in Figure 2E.

**Figure 2:**
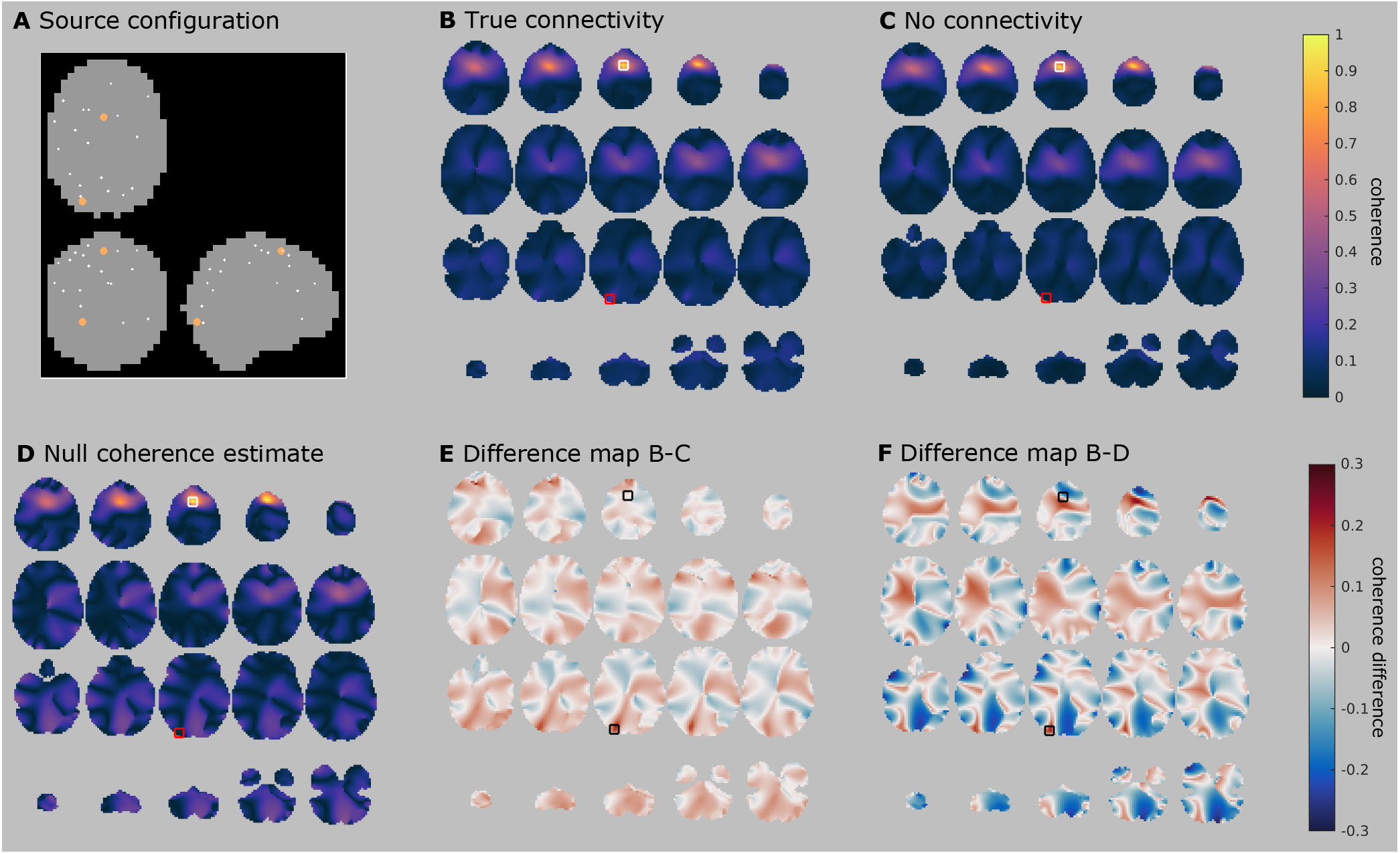
Illustrative example. **A** Spatial configuration for simulation. Shown are 20 randomly selected dipole locations of active sources (small white dots) and the two truly interacting sources (bigger orange dots). **B** Estimated coherence for true connectivity using a single dipole beamformer. The white square denotes the seed, coinciding with one of the interacting sources. The red square denotes the location of the connected dipole. **C** Same as B, but with no underlying interaction. **D** Estimate of null coherence for the same data. **E** Difference map of B and C, black squares denote the interacting dipoles. **F** Difference map of B and D.

Before exploring the usage of different beamformer analysis schemes to improve the connectivity results from Figure 2, let us note that Figure 2 considered a situation in which the seed dipole for the connectivity estimation was well chosen, *i*.*e*., it coincided roughly with one of the truly interacting sources. In the analysis of experimental data, seed locations are not known a priori, thus one might happen to choose locations that are not truly interacting. In this case, the high spatial structure in the null coherence maps is replicated even for non-interacting seeds, which evidently would be problematic for real data analysis. This effect is illustrated in supplementary Figure S1.

### 3.2 Two-dipole beamformer and array subsampling

At this point, one may argue that the suggested null coherence estimate is impractical to use, given the large amount of residual noise in the difference images (*cf*. Figures 1 and S1). In other words, the spurious connectivity estimated between two locations is poorly approximated just by computing the spatial leakage of projected spatially white sensor noise, at least when using a single dipole beamformer formulation. As motivated in the introduction section, the use of a two dipole-constraint in the beamformer formulation may reduce some of the leakage terms in equation 6, leading to a null coherence estimate that is better behaved. In addition, sensor array subsampling allows for multiple (although possibly degraded) estimates of the true structure in the data, while unstructured noise is averaged out when aggregating those estimates. Let us further investigate if the scaled spatial filter inner product might be an appropriate estimate for spurious source interactions when using those alternative beamformer approaches. Figure 3A revisits the results from Figure 2F, plotting the estimated null coherence (x-axis) against the estimated coherence (y-axis) for all dipoles in the stimulation with the values for the interacting dipole pair highlighted with the yellow square. Figure 3B and C show the results for the same single dipole beamformer approach but with the other truly interacting source and a source between the two truly interacting sources as seeds, respectively. Ideally, for non-interacting dipoles, the data points should cluster on a line around the diagonal, while the data point(s) corresponding to the truly interacting dipoles should be clearly above the diagonal. Comparing the single dipole beamformer (Figure 3A and B) with the two-dipole beamformer (Figure 3D and E) for the truly interacting dipoles suggests that, overall, the data points cluster more nicely around the diagonal line in the two-dipole beamformer case. Figure 3G-I depict the results of the subsampling approach. To this end, the average across subsamples of the estimated coherence and null coherence was normalized with the standard deviation of their difference. Here, the subsampling boosts the detectability of the interacting dipole pair, by making it stand out clearly from all other dipole pairs. Also, for seed dipoles in inactive and non-interacting locations (bottom row), the spread of the data points around the diagonal is much more comparable across the different seed dipoles for the subsampling-based reconstruction. In contrast, when no subsampling is used, the deviations from the diagonal are substantially larger for the inactive seed dipole as compared to the active and interacting seed dipoles. This suggests that the magnitude of the spatial noise in the difference images varies considerably, depending on the choice of the seed dipole, and that the approach of array subsampling mitigates this effect by aggregating the results of many different noise realizations.

**Figure 3:**
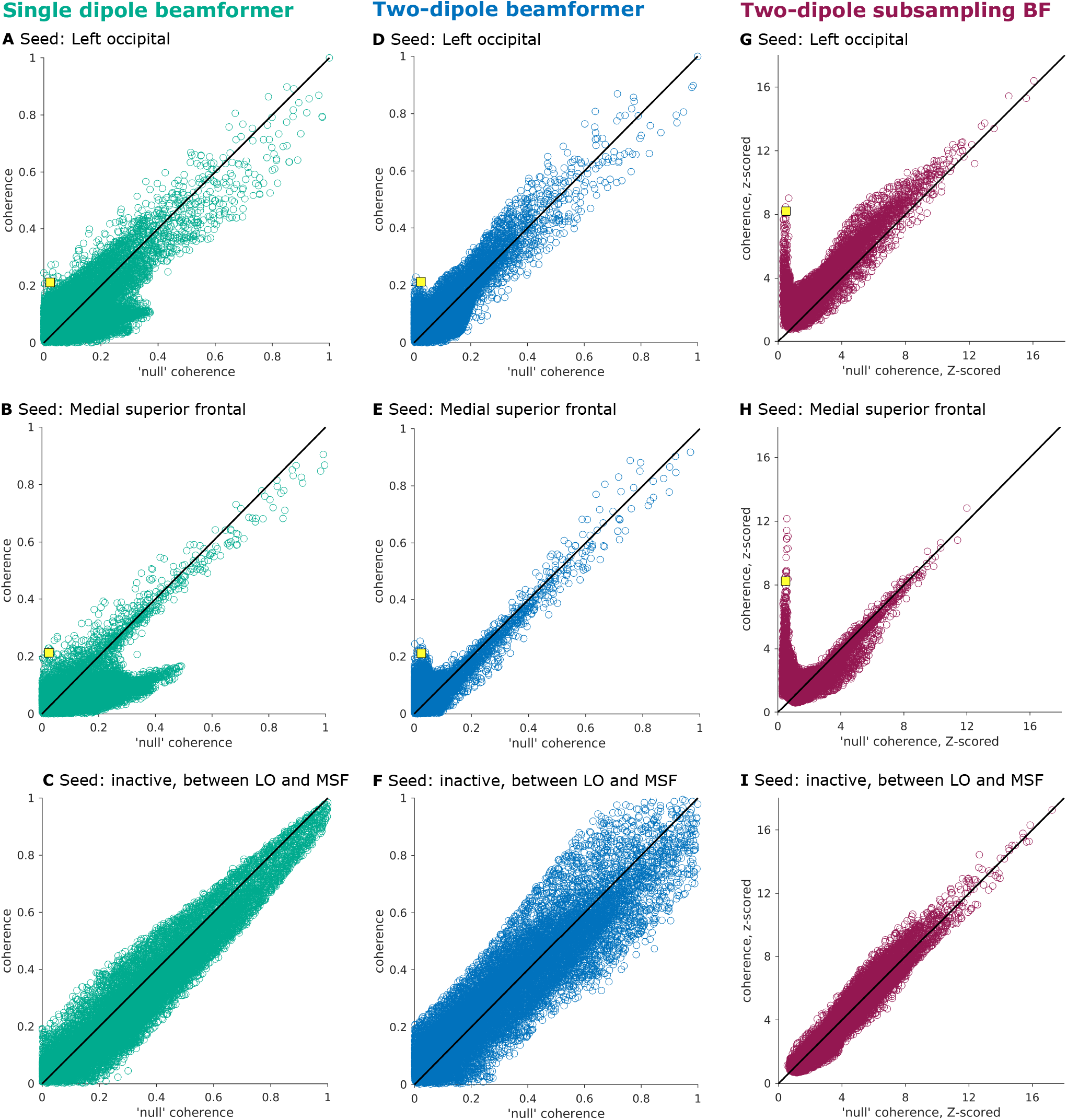
Comparing different beamformer approaches. **A-C** Single dipole beamformer with **A** seed close to truly interacting source in left occipital cortex (LO), **B** seed close to truly interacting source in medial superior frontal cortex (MSF), and **C** seed in a non-active dipole located on the line between interacting dipoles LO and MSF. **D-F** Two-dipole beamformer. **G-I** Two-dipole beamformer with array subsampling. The values for the interacting dipole pair highlighted with the yellow square.

### 3.3 Evaluating all-to-all pairwise coherence

To formally evaluate how the spurious spatial structure in the seed-based connectivity maps interacts with accurate detectability of the true interactions, we constructed and evaluated the all-to-all pairwise coherence matrix (Schoffelen and Gross, 2011). Here, each of the dipoles in the grid serves as a seed dipole to all other dipoles. After the subtraction of an estimate of the null coherence, the resulting 6-dimensional volume of difference in coherence is thresholded, using a relative threshold keeping the *N* % largest values. We explored the following values of N, with the corresponding number of unique supra threshold edges in parentheses: 5% (9.8 *×* 10^5^), 1% (1.95 *×* 10^5^), 0.5% (9.8 *×* 10^4^), 0.1% (1.95 *×* 10^4^), 0.05% (9.8 *×* 10^3^), 0.01% (1.95 *×* 10^3^), 0.005% (975), 0.001% (195), 0.0005% (98).

The thresholded maps are subsequently analyzed for the presence of clusters of spatially connected dipoles in 6-dimensional space. Such clusters are considered to reflect a potential long-distance interaction if they consist of two dipole assemblies that are spatially distinct from each other. Clusters that contain auto-connections, *i*.*e*., dipoles that are present in both assemblies of the connection, are discarded from further inspection. If the simulated interacting dipoles fall within the identified clusters, it is considered a hit. All remaining clusters are considered false positives. It should be noted that the number of false positives evidently will increase with a decreasing threshold when using a relative thresholding scheme as we do here. The total number of false positives further depends on the blurriness of the spatial noise and the degree of auto-connectedness in the data.

Figure 4 shows the clusters with the smallest distance to the simulated interacting dipole pair, and the number of distinct connections, for each of the different thresholds applied. Using a single dipole beamformer (Fig. 4A), the true connection can be correctly identified in three out of the nine thresholding schemes (marked with a red frame). This, however, comes at the expense of additional false positive connections, ranging in number from 70 to 128. Thus, in this relatively favorable context – where coherence is large and the phase of the interaction is close to 90 degrees, *i*.*e*., with only a minor instantaneous correlation between the two sources without the potential corresponding distortion of the beamformer due to correlated sources – the actual connection may be correctly identified, but one has to be prepared to accept an additional large number of false positives.

**Figure 4:**
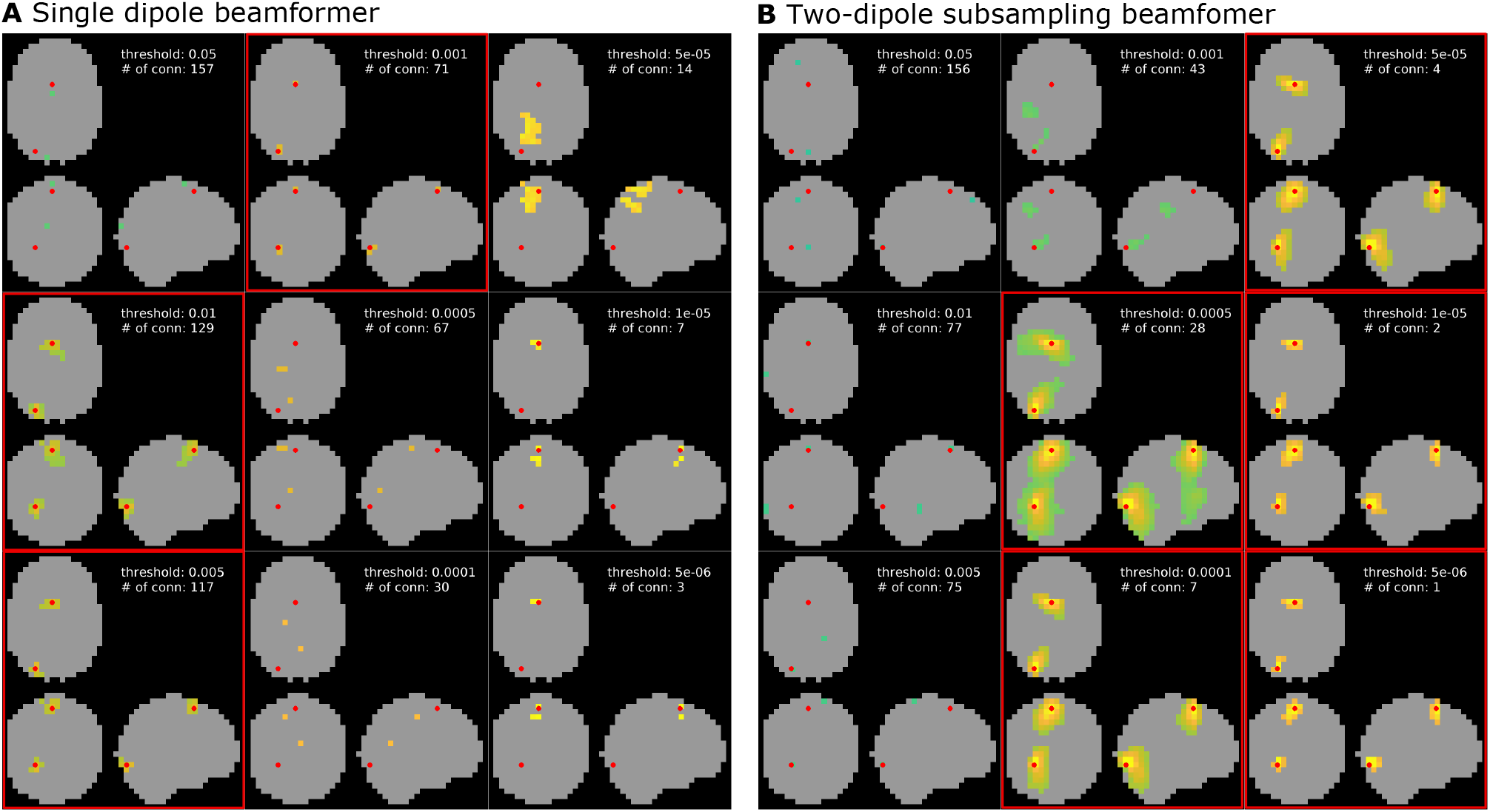
All-to-all pairwise coherence. Shown are the results at different cluster thresholds for the single dipole beamformer (**A**) and the two-dipole beamformer with array subsampling (**B**). Each result also lists the number of identified connections. Thresholds at which the truly interacting dipole pair was successfully identified are marked by a red frame.

Figure 4B shows the spatial clusters closest to the interacting dipole pair for the subsampling-based reconstruction. Here, the interacting sources are correctly identified in the five highest thresholding schemes (marked with red frames), with a considerable reduction in the number of false positives as compared to the single dipole beamformer output in Figure 4A. The number of false positives drops to one or none for the highest two thresholds applied. As an alternative to analyzing the difference in coherence with an approximation of the estimated bias under the assumption of no coherence, one can also investigate the magnitude of the imaginary part of the reconstructed coherency. Figure S2 in the supplementary material replicates the results from Figure 2A (using a single dipole beamformer) for the imaginary part of coherency. With increasing threshold, the true connection can still be reliably identified and the number of false positives drops to only two for the highest two thresholds tested. Importantly, however, the usefulness of the imaginary part of coherency is limited to situations in which the phase difference of the interaction is pointing away from 0 or 180 degrees. Figure 5 shows the results for the same interacting dipole pair as in all previous examples, which are now interacting at a phase difference of zero. The subsampling approach (Fig. 5B) is still capable of detecting the interacting dipole pair at high thresholds, whereas the imaginary part of coherency approach (Fig. 5A) now fails at higher thresholds. Therefore, the subsampling approach with a two-dipole beamformer seems to work well regardless of the phase difference of the interacting dipole pair.

**Figure 5:**
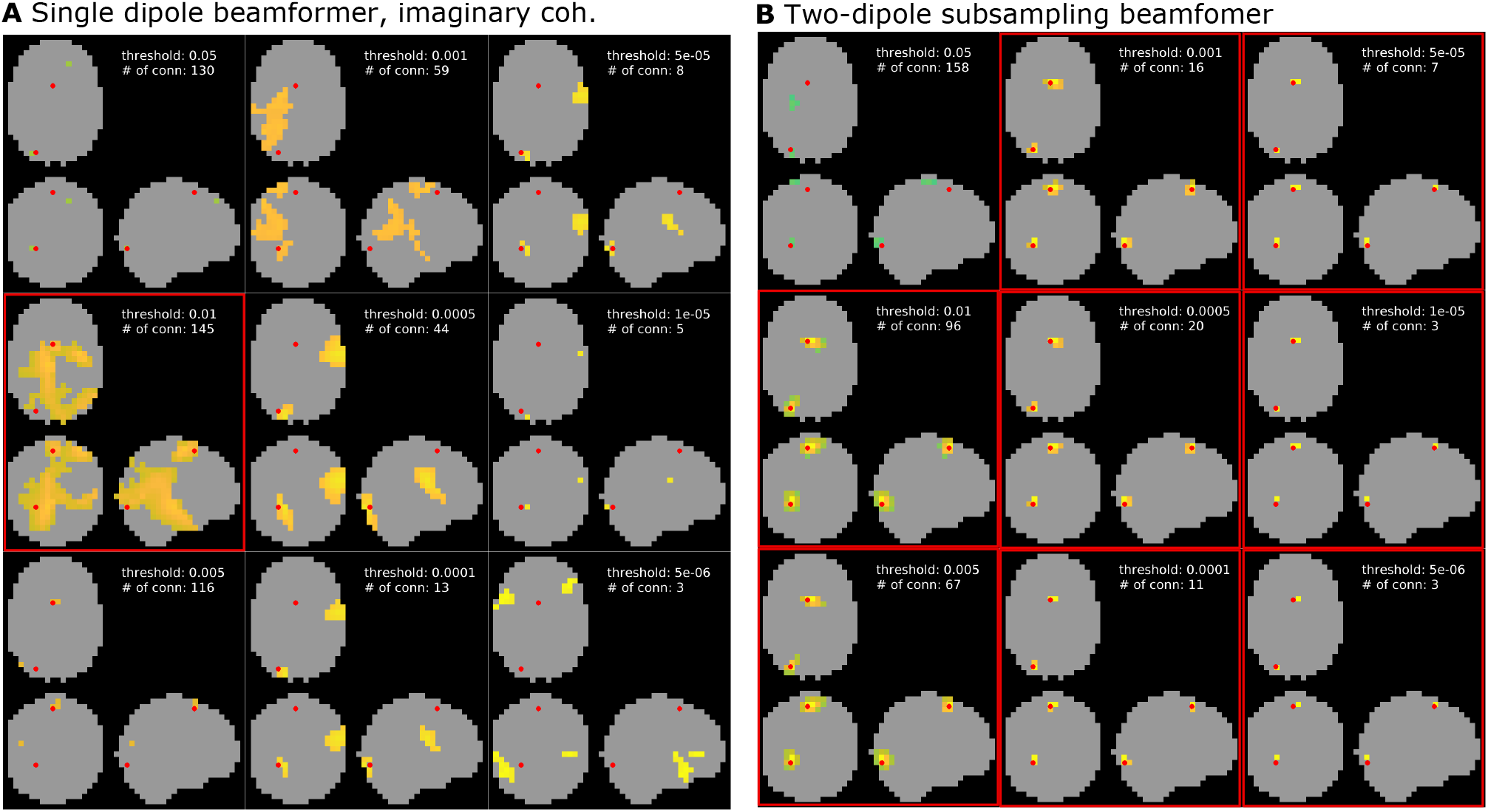
Impact of 0 degree phase shift. Results for interacting sources where the phase of the interaction is 0 degrees for the beamformer with the geometric correction scheme, which focuses on the imaginary part of coherency (**A**) and the two-dipole beamformer with array subsampling (**B**). Each result also lists the number of identified connections. Thresholds at which the truly interacting dipole pair was successfully identified are marked by a red frame.

### 3.4 Full simulation results

To test our proposed approach more thoroughly and to substantiate the illustrative results discussed so far, we employed an exhaustive simulation. Here, we compare the array subsampling two-dipole beamformer approach to three other approaches: the traditional single dipole beamformer, the two-dipole beamformer without subsampling, and a beamformer without sub-sampling, using a geometric correction scheme, proposed by Wens et al. (2015). This correction scheme uses a spatial projection heuristic to remove instantaneous leakage from a seed location’s estimated activity from all target locations’ estimated activity. In practice, this results in the real-valued component of the interaction between the seed and target dipoles to be suppressed, leading to a purely imaginary-valued coherency value. Therefore, in the below, we refer to this last strategy as the reconstruction of the imaginary part of coherency. We evaluate the source reconstruction results based on hit rate, *i*.*e*., how often the chosen approach correctly identified the true interacting dipole pair. Fig. 6 shows the simulation results for a relative amplitude of *a* = 0.7, thus, the interacting sources were 2.333 times stronger than the other active sources (for the results for *a* = 0.5 and *a* = 0.8, we refer the reader to supplementary Figures S3 and S4, respectively). The results reported in the paper are based on a signal-to-sensor-noise ratio of 0.6, the results for an SNR of 0.5 are reported in supplementary Figures S6-S8.

**Figure 6:**
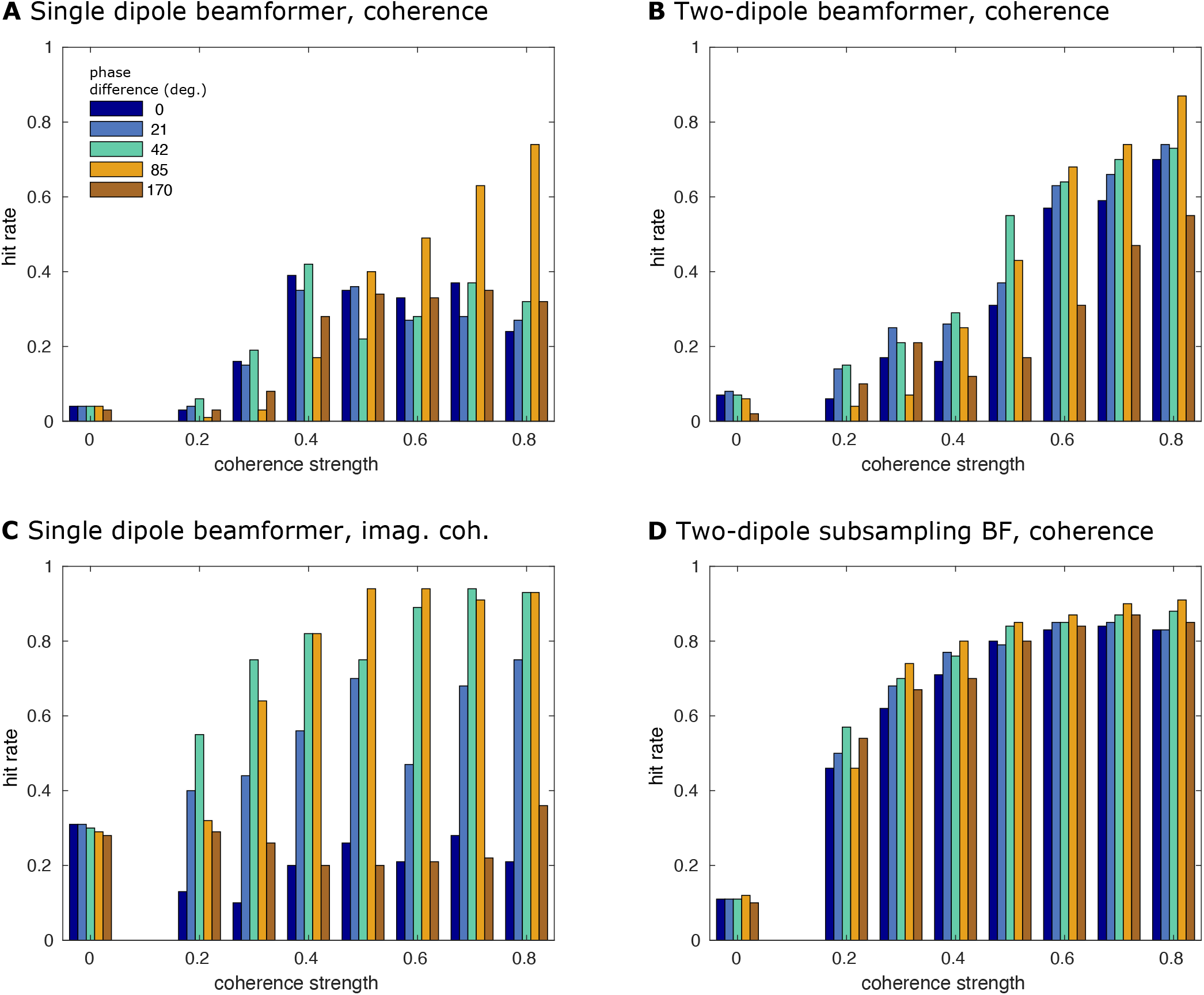
Detection rate for interacting dipole pairs. Results from the full simulation, showing the hit rates for the interacting dipole pair as a function of simulated coherence strength and phase difference. The relative amplitude of the interacting sources and the other sources was a = 0.7, *i*.*e*., the interacting sources were 2.333 times stronger than the other active sources. The SNR was 0.6. **A** Traditional single dipole beamformer. **B** Two-dipole beamformer. **C** Single dipole beamformer using imaginary coherence. **D** Two-dipole beamformer with array subsampling.

Fig. 6 depicts the hit rate as a function of simulated coherence strength, and phase difference, for the different reconstruction strategies. We first considered the situation in which application of at least one of the thresholds *<* 0.01% resulted in the detection of the dipole pair that was chosen for the interaction (to define a hit, we allowed the summed distance of the simulated dipoles to the closest voxel in the suprathreshold clusters to be at most 2 cm). Overall, the performance of the single dipole approach (Fig. 6A) was quite poor, with the hit rate — as a function of coherence strength and phase difference — rarely exceeding 40%. Only at unrealistically high coherence strengths > 0.6 was the detection rate larger than 50%, and even then only at phase differences close to 90 degrees. The two-dipole approach (Fig. 6B) fared better, specifically for coherence values larger than 0.4. The single dipole beamformer using imaginary coherency (Fig. 6C) generally showed better performance, already at lower coherence values, but this performance was highly dependent on the phase difference of the interaction, where interactions with a phase difference close to 90 degrees were more readily detectable, reaching a hit rate of > 90% in some situations. When the phase difference of the interaction was close to 0 (or 180) degrees, however, the detection rate at higher coherence values was only slightly higher than for low coherence strengths, compared with the single and two-dipole approach. The array subsampling beamformer (Fig. 6D) overall performed best. Even though the maximum detection rate was not as high as in some situations using the imaginary part of coherency (*i*.*e*., coherence > 0.5 and phase difference close to 90 degrees), the detection rate at a moderate coherence of 0.3 already exceeded 60%, independent of the phase difference. Thus, the array subsampling two-dipole beamformer outperforms the other approaches for almost all parameters, specifically considering the fact that physiologically realistic neuronal interactions are not constrained to phase differences close to 90 degrees, nor are those interactions restricted to high coherence values. The findings depicted in Fig. 6 are at large supported by the results of other amplitude and SNR values, as supplementary Figures S4 and S6-S8 illustrate.

Notably, in the absence of simulated true coherence, the different reconstruction approaches resulted in a variable amount of false positive connections in the direct vicinity of a pair of activated dipoles (see the leftmost set of bars in each of the panels in Fig. 6). For the imaginary part of coherency this type of false positive connection was present in about 30% of the simulations, and for the proposed subsampling approach the percentage of occurrence was about 10%. In general, the occurrence of false positives is the consequence of the fact that we used a relative thresholding scheme to investigate the spatial structure of the reconstructed connectivity maps. By construction, and irrespective of the numeric value of the connectivity estimates, the relative thresholding scheme always results in a collection of suprathreshold edges in the connectivity maps, which may be spatially clustered, and interpreted as interacting sources. Based on the spatial smoothness of the connectivity maps, and the number of suprathreshold edges, the number of false positive connections will vary as a function of the chosen threshold. Fig. 7 shows the number of false positives versus the hit rate in a so-called Free-response Receiver-Operating-Characteristic (FROC), as a function of the detection threshold and for a relative amplitude of *a* = 0.7. On each of the lines, the threshold is increasing from left to right. For all but the subsampling reconstruction method, the optimal — yet still quite low — sensitivity was reached at a threshold that yielded close to 100 false positive connections on average. For the subsampling reconstruction method, the highest sensitivity was compromised by about 10 to 20 false positive connections on average. Although this still may seem a rather high false positive rate, it is substantially lower than the false positive rate for the other approaches tested. The FROC curves for relative amplitudes of *a* = 0.5 and *a* = 0.8 can be found in the supplementary material (Fig. S5) and show very similar patterns. For an SNR of 0.5, the results are reported in supplementary Figure S9 and at large support the findings for an SNR of 0.6, except for at a low relative amplitude of *a* = 0.5, the only parameter combination for which the two-dipole subsampling beamformer does not clearly outperform the other algorithms.

**Figure 7:**
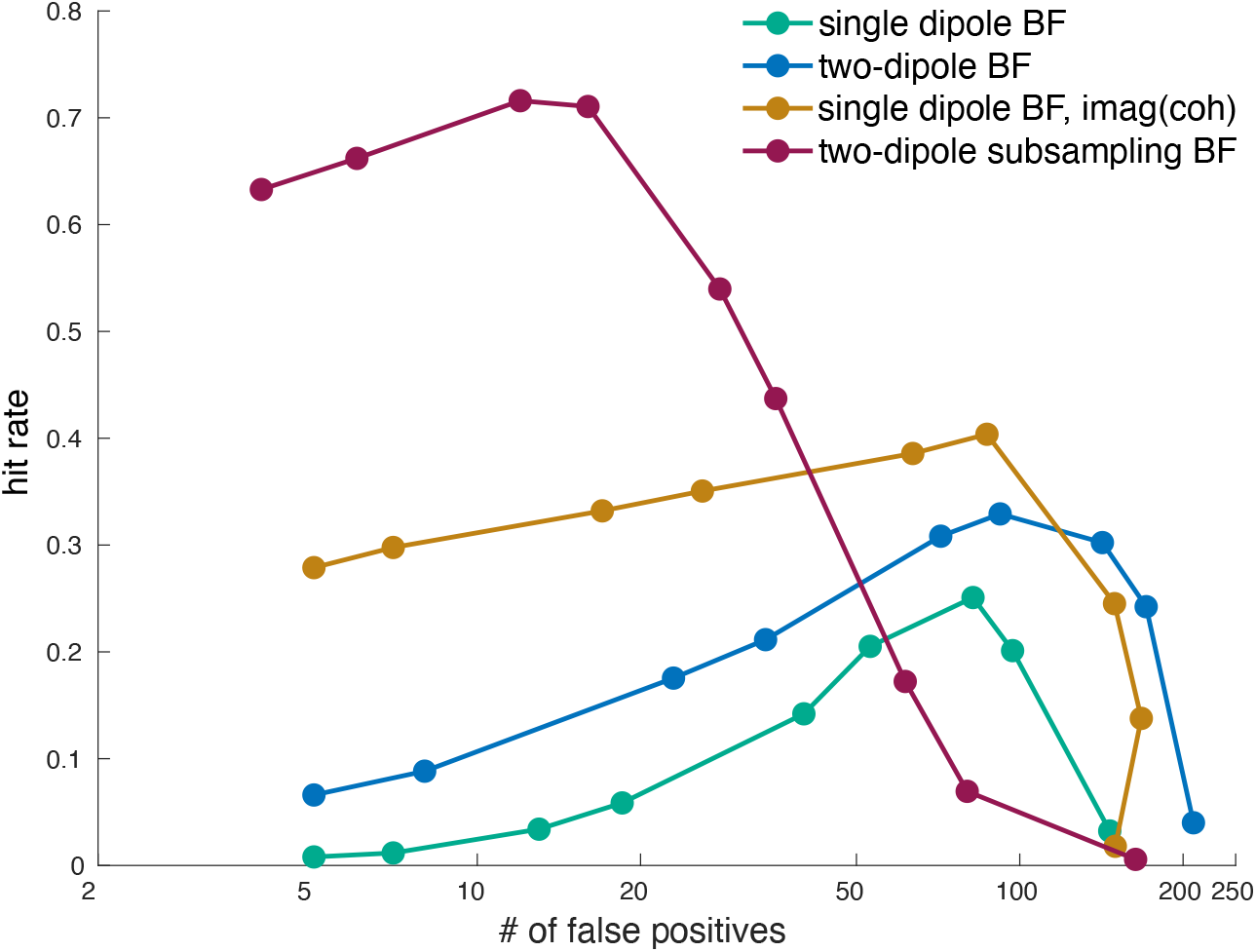
Free-response Receiver-Operating-Characteristic. Hit rate plotted against the number of false positive connections at a relative amplitude of *a* = 0.7.

## 4 Discussion and future directions

Brain connectivity plays a central role in many prevalent hypotheses on brain functioning and organization (Fries, 2005; Jensen and Mazaheri, 2010; Fries, 2015; Bonnefond et al., 2017). Thus, the estimation of functional connectivity based on electrophysiological processes is a necessary tool for the experimental assessment of those theories. Over the years, many different measures of brain connectivity have been put forward (Aviyente et al., 2011; Vinck et al., 2011; Ghanbari and Moradi, 2020) and the methods to apply these have been refined (Dalal et al., 2006; Kuznetsova et al., 2021; Woolrich et al., 2011; Hillebrand et al., 2012; Nunes et al., 2020). Despite these efforts, results from non-invasive recordings, especially phase synchrony measures, have stayed sparse and methodological challenges remain (Colclough et al., 2016; Bastos and Schoffelen, 2016; Palva et al., 2018; He et al., 2019; Schoffelen and Gross, 2009). In this paper, we aimed at addressing some of these challenges through a new beamformer-based connectivity estimation framework, which utilizes three key components: a two-dipole beamformer approach to estimate all-to-all connectivity, an estimation of the null coherence of the model under the assumption of no interaction, and a sensor array subsampling approach to further mitigate the influence of spatial noise. Our all-to-all approach is motivated by the fact that — even in an unrealistically well-controlled contrast — misspecification of the seed dipoles leads to spatial structure in connectivity difference maps that can be mistaken for true interactions. Moreover, because experimental contrasts almost invariably contain differences in source activations and SNR, difference maps of connectivity may show spatial structure that is not due to changes in actual interaction between sources. For this reason, it is desirable to estimate the spatial leakage of connectivity directly from the data. We explored the possibility to use such null coherence estimates, based on the weighted inner product between pairs of spatial filters. A two-dipole beamformer model is motivated by the notion that beamformer estimates are distorted in the presence of underlying correlations. Furthermore, we propose to use sensor array subsampling in order to smooth out the spatial noise at the benefit of the true interactions.

Some of the key components of our approach have been proposed before, in one form or another, but mostly with a different intention, and were never combined for the assessment of connectivity. We compared the performance of our approach to other all-to-all reconstruction schemes, which used only a subset — or none — of the key components.

Using an extensive set of simulations we showed that our approach outperforms the other, often more traditional, all-to-all approaches tested. The overall detection rate, specifically at physiologically meaningful interaction strengths and at a wide range of phase angles, was highest for the proposed subsampling based method. This high detection rate was accompanied by the overall lowest false positive rate. While performance was considerably affected when the relative source amplitude of competing, non-interacting sources was increased (supplementary Figure S3), our approach still showed the overall lowest false positive rate (Figure S5A). Based on these observations, we argue that the proposed reconstruction approach can be a promising pipeline to be evaluated on real MEG data for the robust detection of phase synchronization in brain networks.

Apart from the evaluation of the utility of the proposed approach on real experimental data, we foresee future work to explore in more detail certain aspects of the proposed analysis scheme. For instance, regarding the subsampling, we have settled on a fixed number of subsampled reconstructions, using a random number of sensors (between 50 and 150), and combined the reconstructions by means of averaging. Although those parameter choices were motivated by initial explorations, strategies to estimate the optimal number of sensors for the subsampling, and different combinatorial strategies (*e*.*g*., by also taking the variance structure across subsample based reconstructions into account) may further improve the performance of the subsampling based approach.

## Acknowledgments

This work was supported by the European Union’s Horizon 2020 Marie Sk-lodowska Curie Individual Fellowship (grant agreement 893912) to BUW.

