## Supplementary Material for "Towards a more robust non-invasive assessment of functional connectivity"

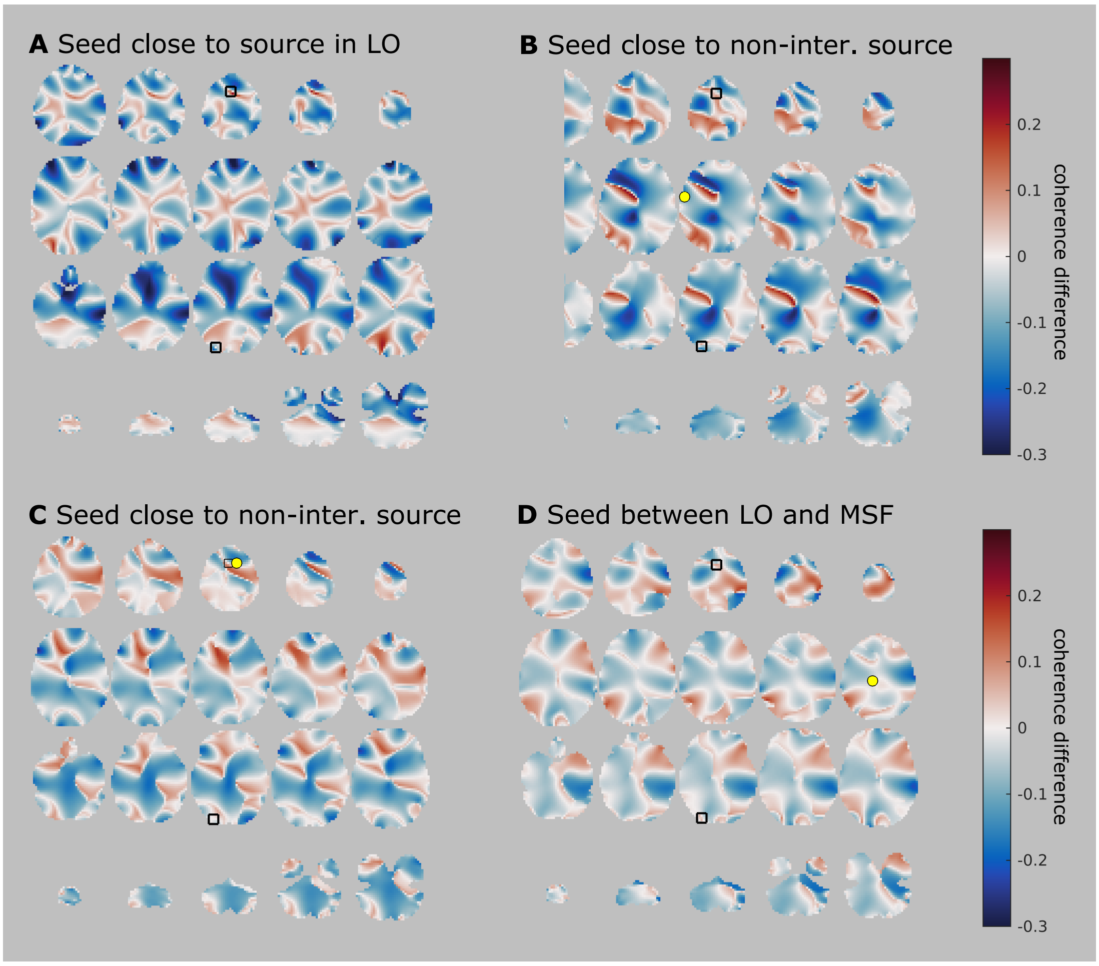


Figure S1: The figure shows the coherence difference maps (using the null coherence estimate) for four other seed locations. **A** The seed dipole almost coincides with the LO source, and thus would ideally show a map that at least shows a local (and ideally global) maximum at the location of the interacting source. **B, C and D** reflect seed dipoles at the location of the yellow dots and should ideally show low coherence difference values without clear local maxima. The seed locations in **B and C** were close to active (but non-interacting) sources, and the seed location in **D** was in the middle of the line segment connecting the two interacting sources. Compared to Figure 1, it is clear that there is ample spatial structure in those maps, even when a seed is placed close to a non-interacting active source or even at a silent location. This is problematic in real applications, because conclusions with respect to interacting sources are typically based on the identification of local maxima in (thresholded) seed-based connectivity (difference) maps.

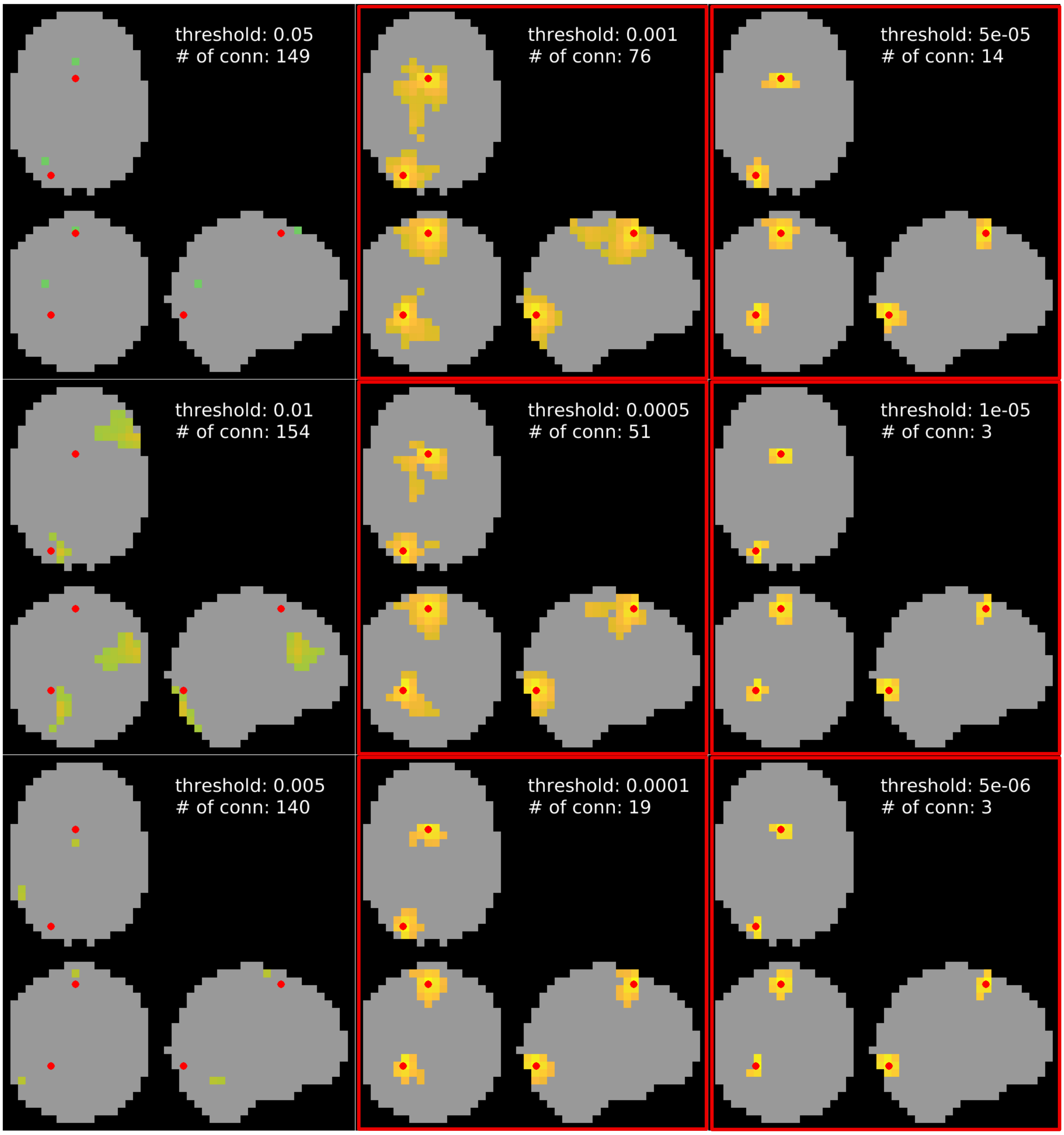


Figure S2: Results from a beamformer with geometric correction, which focuses on the imaginary part of coherency. Each result also lists the number of identified connections. Thresholds at which the truly interacting dipole pair was successfully identified are marked by a red frame.

### Full simulation results for signal-to-noise ratio of 0.6

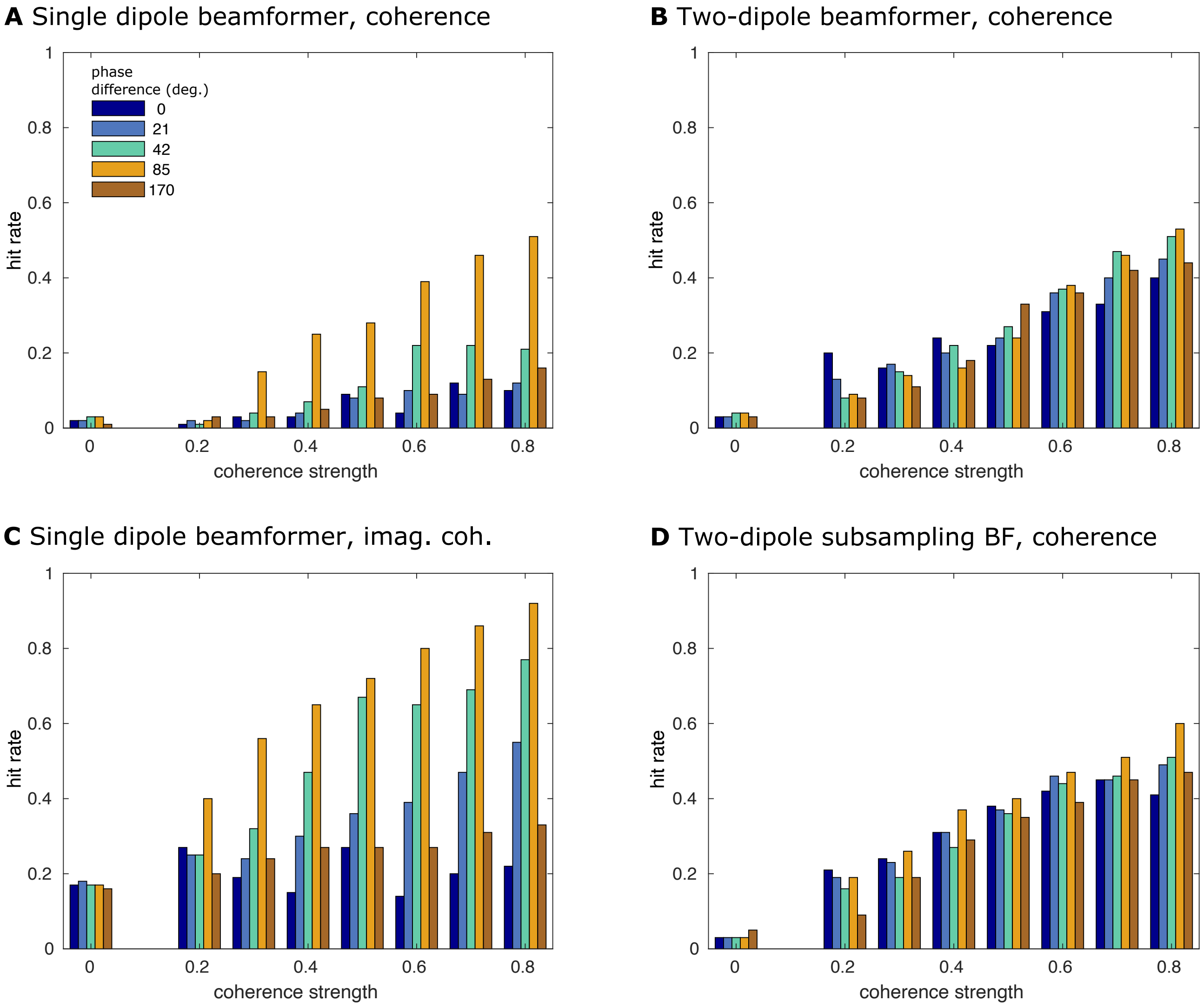


Figure S3: **Detection rate for interacting dipole pairs, a = 0.5.** Results from the full simulation, showing the hit rates for the interacting dipole pair as a function of simulated coherence strength and phase difference. The relative amplitude of the interacting sources and the other sources was a = 0.5, *i.e.*, the interacting sources were equally strong as the other active sources. **A** Traditional single dipole beamformer. **B** Two-dipole beamformer. **C** Single dipole beamformer using imaginary coherence. **D** Two-dipole beamformer with array subsampling.

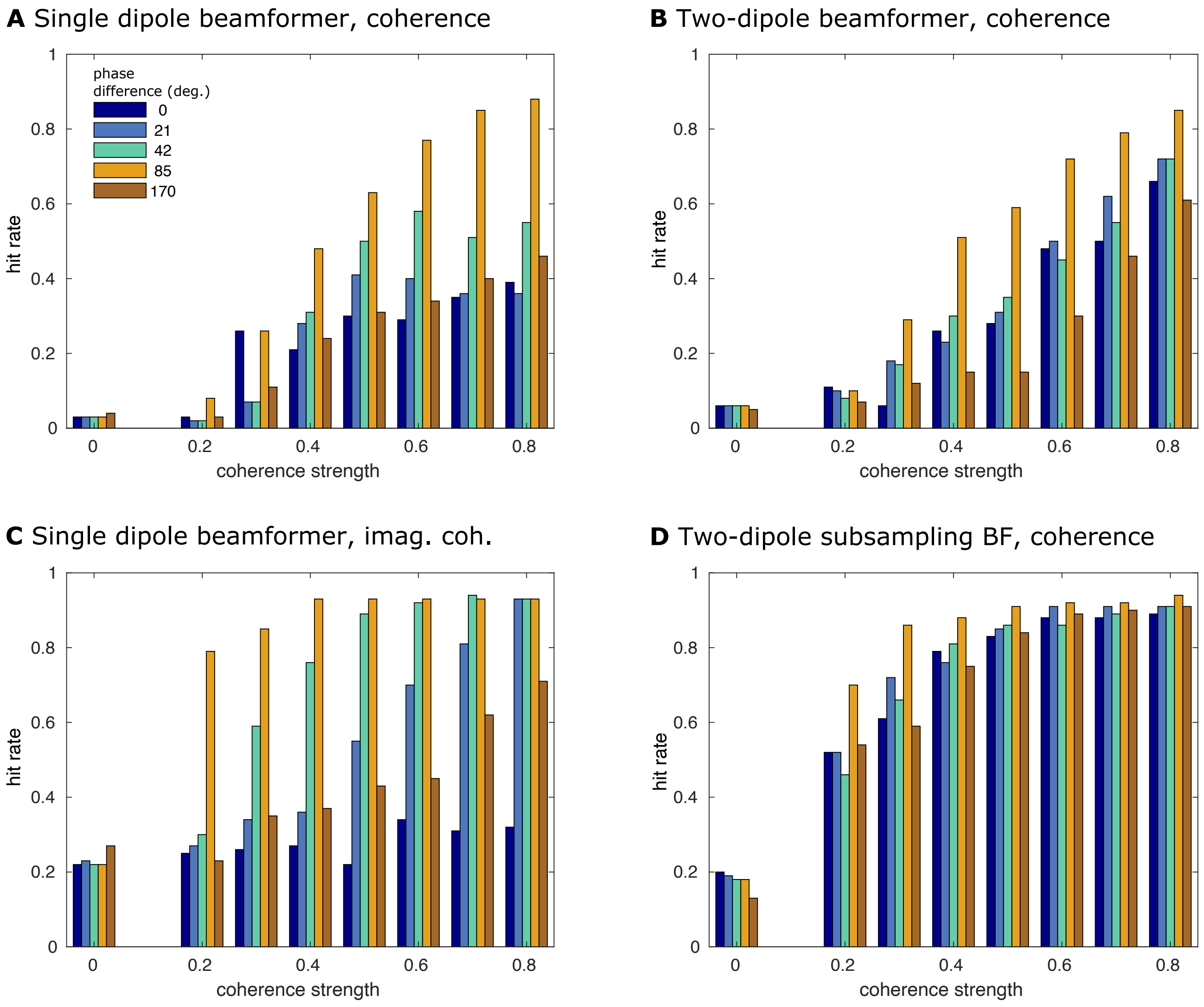


Figure S4: **Detection rate for interacting dipole pairs, SNR = 0.6 and $a$ = 0.8.** Like Figure S3, but the relative amplitude of the interacting sources and the other sources was $a = 0.8$, *i.e.*, the interacting sources were 4 times stronger than the other active sources.

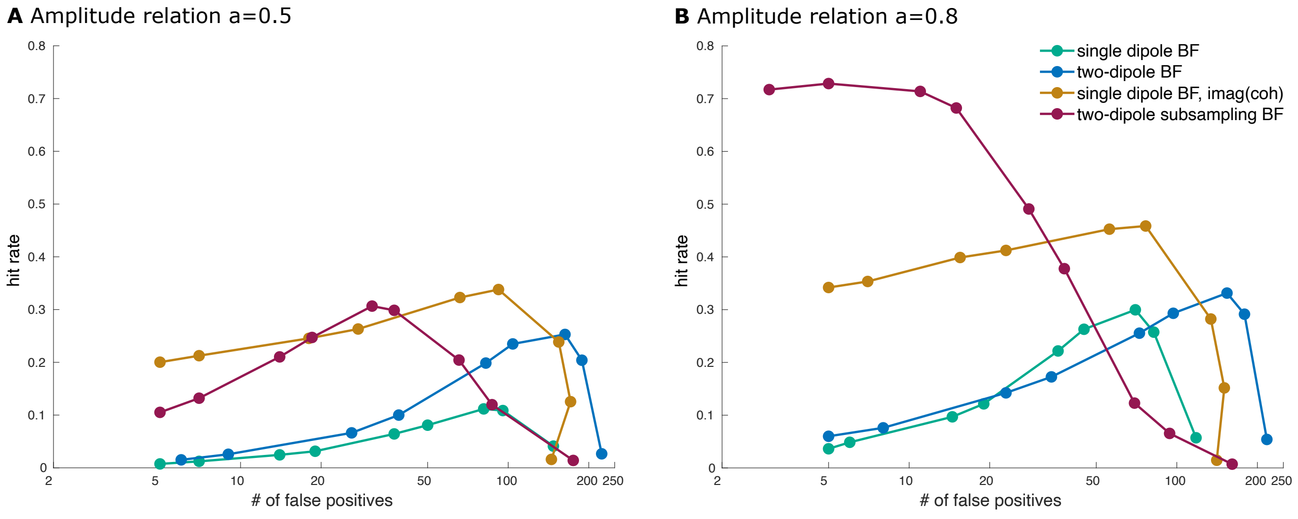


Figure S5: **Free-response Receiver-Operating-Characteristic, SNR = 0.6.** Hit rate plotted against the number of false positive connections at a relative amplitude of **A** $a = 0.5$ and **B** $a = 0.8$.

### Full simulation results for signal-to-noise ratio of 0.5

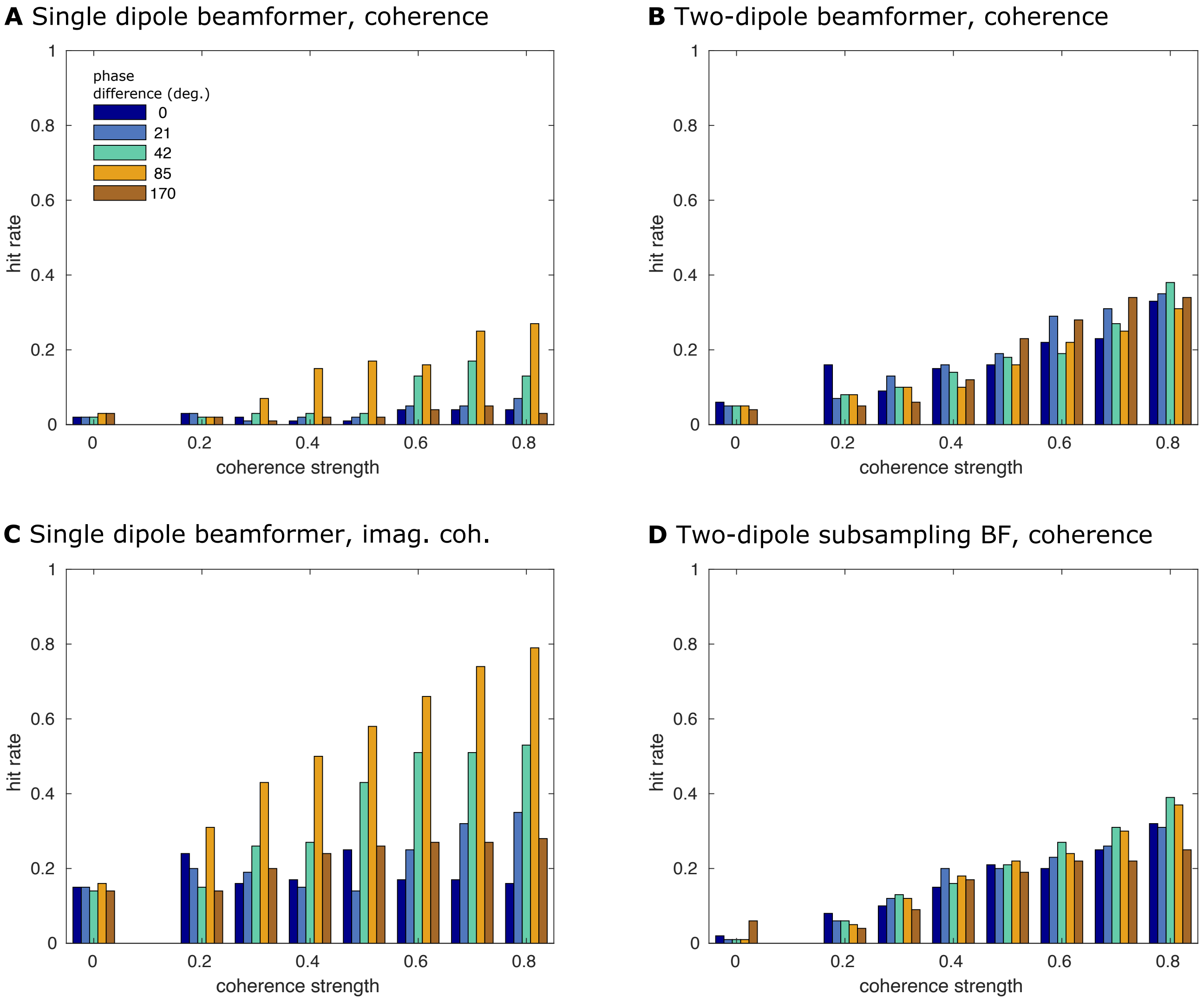


Figure S6: **Detection rate for interacting dipole pairs, SNR = 0.5 and $a$ = 0.5.** Like Figure S3, but for a signal-to-noise ratio of 0.5 instead of 0.6. The relative amplitude of the interacting sources and the other sources was $a = 0.5$.

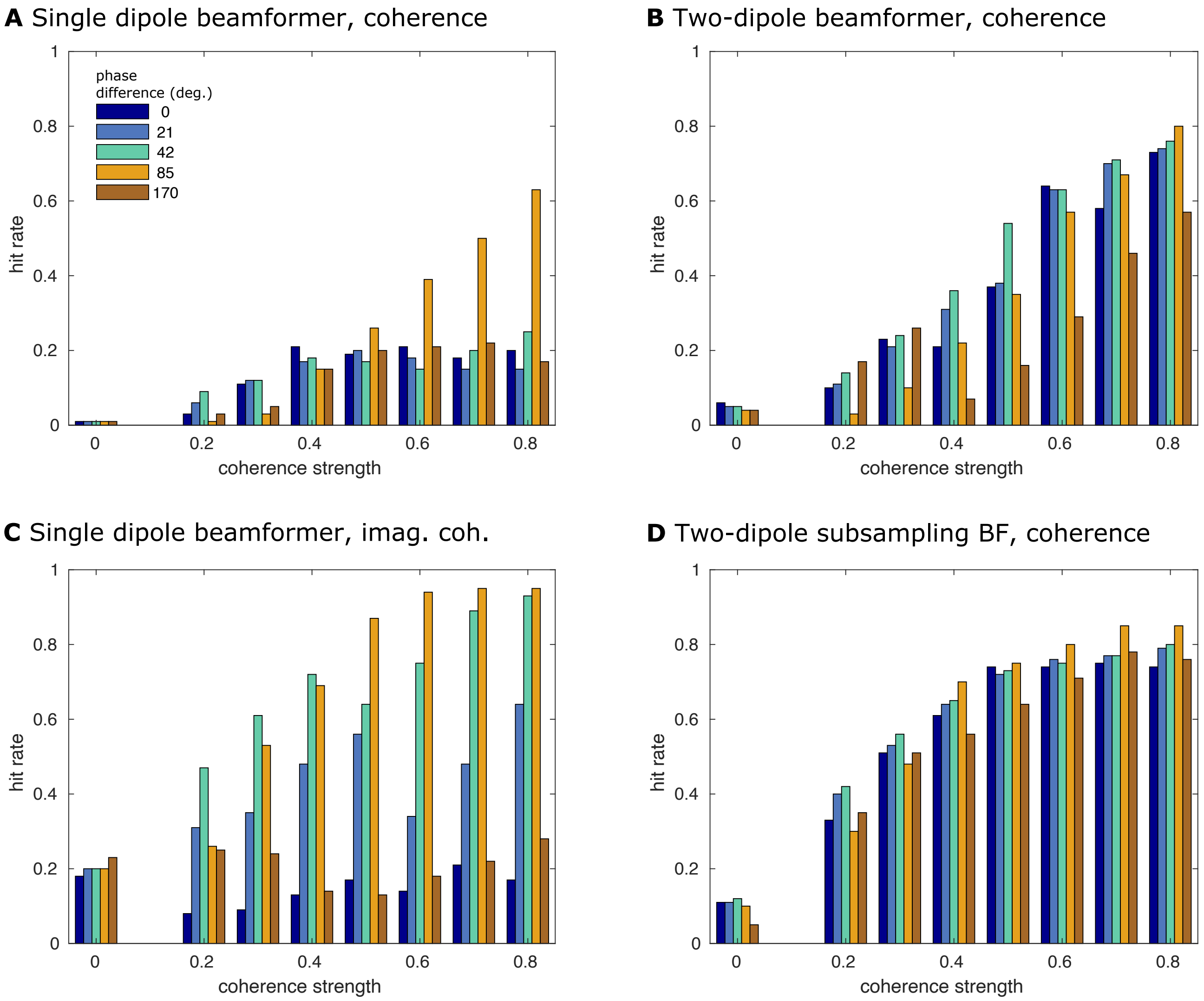


Figure S7: **Detection rate for interacting dipole pairs, SNR = 0.5 and $a$ = 0.7.** Like Figure S3, but for a signal-to-noise ratio of 0.5 instead of 0.6. The relative amplitude of the interacting sources and the other sources was $a = 0.7$.

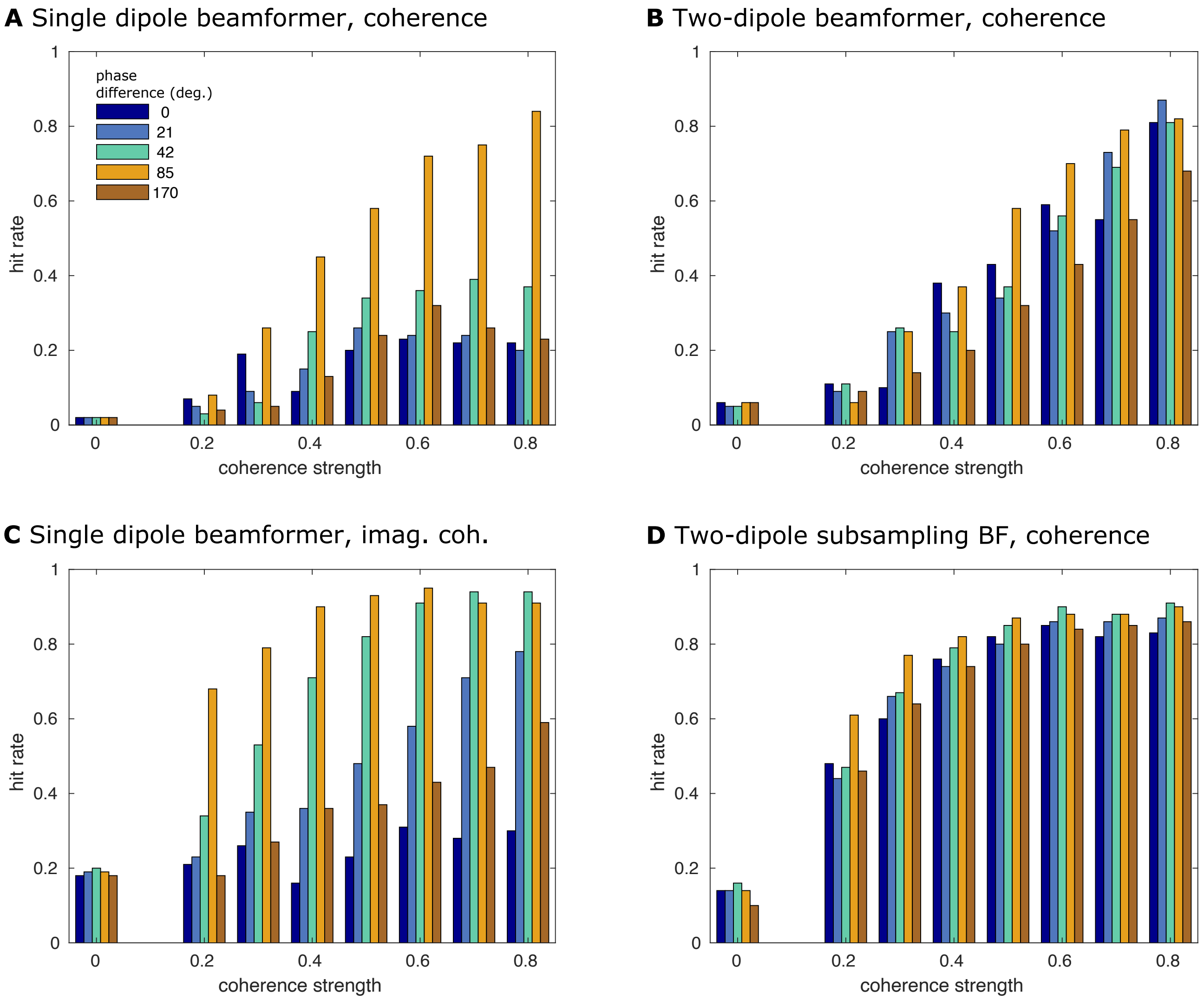


Figure S8: **Detection rate for interacting dipole pairs, SNR = 0.5 and $a$ = 0.8.** Like Figure S3, but for a signal-to-noise ratio of 0.5 instead of 0.6. The relative amplitude of the interacting sources and the other sources was $a = 0.8$.

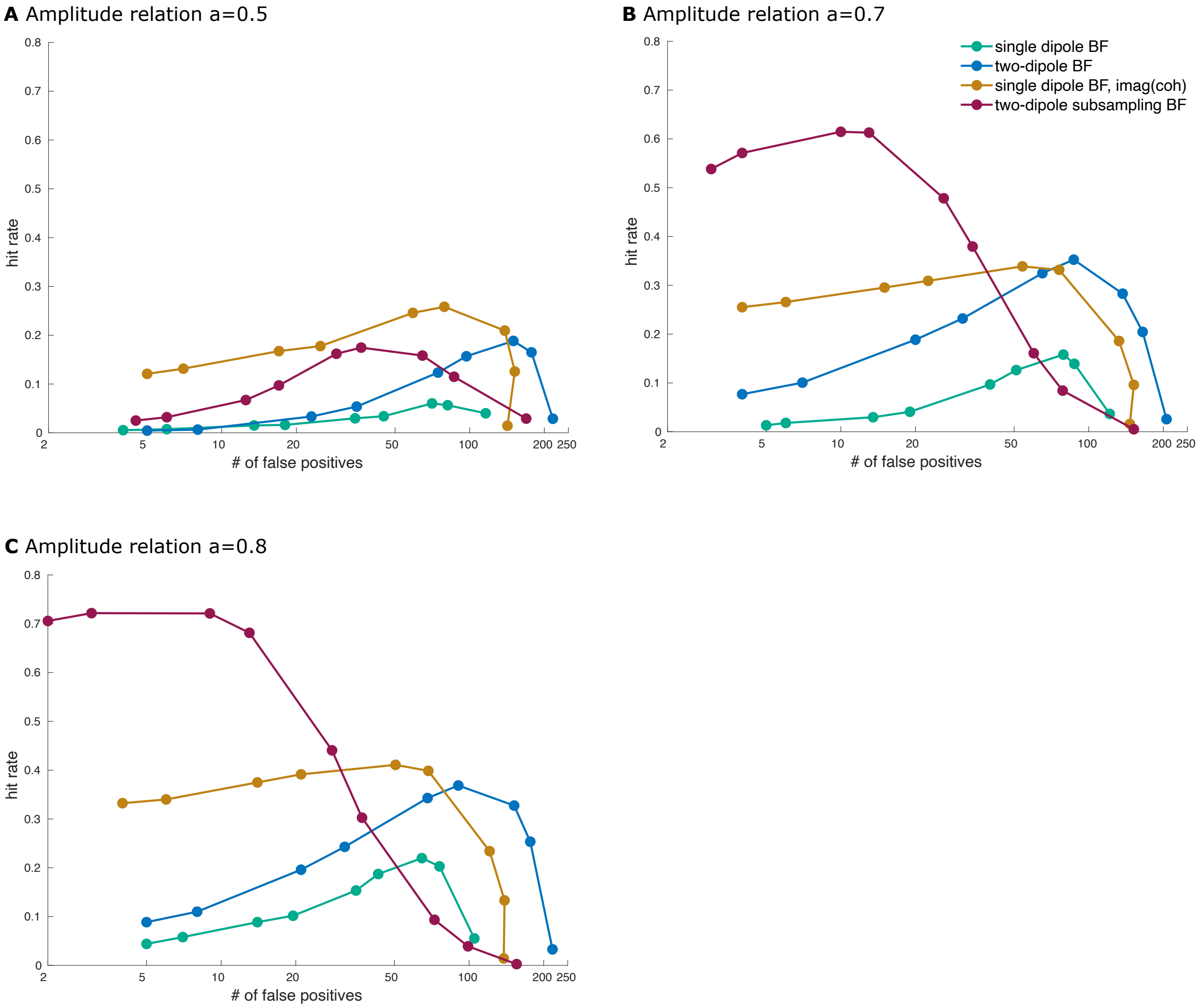


Figure S9: **Free-response Receiver-Operating-Characteristic, SNR = 0.5.** Hit rate plotted against the number of false positive connections at a relative amplitude of **A** $a = 0.5$, **B** $a = 0.7$, and **C** $a = 0.8$.
